# Optimization of rearing *Transeius montdorensis* under laboratory conditions

**DOI:** 10.1101/2024.09.13.612991

**Authors:** Hung Nguyen, Binh Nguyen, Bishwo Mainali, Maciej Maselko

## Abstract

The global application of *Transeius montdorensis* (Acari: Phytoseiidae) as a biological control agent across various protected crops has proven effective against a range of insect pests like thrips and whiteflies, as well as pest mites like broad mites and russet mites. Optimization of rearing *T. montdorensis* under laboratory conditions is crucial for further studies of this species to improve their application in Integrated Pest Management (IPM) programs. Here, we evaluated the development and reproduction of *T. montdorensis* when fed on four different diets, including cattail pollen (*Typha latifolia*), living dried fruit mites (*Carpoglyphus lactis*), frozen *C. lactis* eggs, and a mixed diet of frozen *C. lactis* eggs and *T. latifolia* pollen. Females consuming the mixed diet exhibited superior total fecundity and daily oviposition rate, along with the highest intrinsic rate of increase (r_m_) and net productive rate (R_0_) among all diets tested. The immature period was significantly longer for mites on a diet of living *C. lactis* compared to those on other diets. Importantly, utilizing frozen *C. lactis* eggs and *T. latifolia* pollen mitigates the risk of infestation and contamination from the living dried fruit mites, which is important for laboratory and field settings when releasing the predator colonies. Our findings not only present an optimized rearing method for predatory mites under laboratory conditions but also suggest potential broader applications for enhancing the effectiveness and sustainability of biological control strategies across various agroecosystems and reducing dependency on chemical pesticides.

## Introduction

Within the group of phytophagous predatory arthropods that feed on both plants and animals, predacious mites of the Phytoseiidae family play a vital role in controlling key agricultural pests, including thrips, whiteflies, and phytophagous mites, which negatively affect economically important crops (Bazgir et al., 2019; Chant, 1985; Samaras et al., 2019). Demite et al. (2014) reported that the Phytoseiidae family comprises approximately 2,709 formally recognized species classified into 91 distinct genera. Based on their feeding habits, biological traits, and morphological features, those predatory mites were categorized into four primary types (with several sub-types): specialized mite predators of Tetranychus species, selective predators of tetranychid mites, generalist predators, and polliniphagous generalist predators (JA McMurtry & Croft, 1997). This family stands out as the initial and primary group of predatory mites that colonize foliage environments in various ecosystems (McMurtry, 2010). Their increasing incorporation into Integrated Pest Management (IPM) programs across different cultivated crops highlights their crucial role in reducing reliance on chemical insecticides, offering a sustainable approach to managing agricultural insect pests (Javier et al., 2010; Van Lenteren, 2012).

*Transeius montdorensis* (formerly known as *Amblyseius montdorensis/ Typhlodromips montdorensis/ Typhlodromus montdorensis*) (Acari: Phytoseiidae), a polyphagous predatory mite, was originally found in New Caledonia and was first described by Schicha (Schicha, 1979; Steiner et al., 2003). According to their food preference, *T. montdorensis* was categorized into type III, generalist predators (JA McMurtry & Croft, 1997). This predator targets a variety of pests, including *Frankliniella occidentalis* (Labbé et al., 2019; Rahman et al., 2011), whiteflies (Cuthbertson, 2014; Sun et al., 2022), spider mites (Schicha, 1979), and broad mites (Steiner & Goodwin, 2002). Though originating from the subtropical areas of Australia, *T. montdorensis* thrives in temperatures between 20 °C and 30 °C (Cox et al., 2006) and has adapted to colder environments, with a developmental threshold between 10.3 °C and 10.7 °C observed in the UK (Hatherly et al., 2004). Due to their wide range of prey, high adaptation in different environments as well as the potential to sustain their population at relatively high levels even at low prey or in the absence of prey (JA McMurtry & Croft, 1997; McMurtry et al., 2013), *T. montdorensis* has been positioned as a commercial beneficial agent for pest control in global agricultural settings, being available in Australian and European markets since 2003 (Van Lenteren, 2012). Furthermore, Steiner et al. (2003) and Buitenhuis et al. (2010) indicated that *T. montdorensis* outperforms *Neoseiulus cucumeris* in intrinsic rate of increase, daily oviposition rate, and thrips larvae consumption. Another study also reported that *T. montdorensis* is more effective than *Amblydromalus limonicus* in controlling two whitefly species, *Bemisia tabaci* and *Trialeurodes vaporariorum*, especially under cold temperatures (Richter, 2016). On the contrary, Mouratidis et al. (2023) concluded that *A. limonicus* and *Ambleyseius swirskii* performed better in controlling *Scirtothrips dorsalis* on strawberries than *T. montdorensis* and *N. cucumeris*. In Australia, *T. montdorensis* has been successful in managing *F. occidentalis* across various crops (Manners et al., 2013; Steiner & Goodwin, 2002), showing a higher intrinsic rate of increase (r_m_) than *N. cucumeris* (Steiner et al., 2003) and superior in daily *F. occidentalis* consumption than *N. cucumeris* on strawberry (Rahman et al., 2012).

Type III predatory mites can develop and reproduce on diverse diets, including natural prey like phytophagous mites, thrips, and whiteflies, as well as alternative food sources like pollen, honeydew, and factitious prey like storage mites (Jaques et al., 2015; Nguyen et al., 2013). Although pollen quality differs among plant species and environmental conditions (Lundgren, 2009), it is favored due to the high quality and abundant of pollen grains, cost-effectiveness, ease of collection, and its suitability in enhancing the development and reproduction of phytoseiid mites (Irina Goleva & Claus PW Zebitz, 2013; Gravandian et al., 2022; Samaras et al., 2015; Vangansbeke et al., 2014). Schreiber (2018) noted that pollen serves as both an alternative and supplementary food source for feeding predatory mites. Interestingly, several predacious mites are capable of growing and reproducing solely on pollen (Ranabhat et al., 2014; Steiner et al., 2003). Among a number of plant pollen species, cattail pollen (*Typha latipholia*) is a commercial product and a promising alternative food source for mass-rearing predatory mites (Gravandian et al., 2022).

Due to the high costs associated with maintaining and rearing generalist predators using natural prey, storage mites have been applied as a cost-effective and less laborious candidate for rearing various predatory mites (JA McMurtry & Croft, 1997; Pirayeshfar et al., 2020). These factitious hosts are employed not only to feed predatory mites under laboratory conditions or in mass-rearing programs but also serve as an additional food source to sustain populations of predatory mites in the crops when their preferred prey is at low density or absent (Barbosa & de Moraes, 2015; Massaro et al., 2021). Moreover, prey mites do not support the development and reproduction of thrips (Pirayeshfar et al., 2020). Among various storage mites, *Carpoglyphus lactis* has been extensively researched and utilized for rearing different generalist predators and has shown promising results (Nguyen, 2015; Wang et al., 2024). However, the required space and labor make them costly (Gravandian et al., 2022). Additionally, their potential to cause infestations and health issues, such as gastrointestinal acariasis, allergic dermatitis, and respiratory allergies in humans, poses significant risks in food production (Hubert et al., 2018; Jan Hubert, 2011; F. Pirayeshfar et al., 2021b).

Alongside the mentioned diets, including plant food and living storage mites, frozen stages of factitious hosts were recognized as a potential diet for feeding several species of predatory mites such as *A. swirskii* (Pirayeshfar et al., 2020) and *Blattisocius mali* (F. Pirayeshfar et al., 2021a). Moreover, a combination of factitious host and pollen has been applied to investigate the developmental and reproductive responses of predacious mites, including *Amblydromalus lomonicus* (Samaras et al., 2019) and *B. mali* (Pirayeshfar et al., 2022). Similarly, another research demonstrated that a mixture of almond pollen and the flour moth *E. kuehniella* eggs significantly enhanced the oviposition of *N. cucumeris* (Etienne et al., 2021). However, no studies have examined the impact of frozen factitious food alone or mixed with pollen on *T. montdorensis*. Therefore, this study assessed life table parameters using four diets, including cattail pollen (*T. latifolia*), living dried fruit mite (*C. lactis*), frozen *C. lactis* eggs, and a mixture of frozen *C. lactis* eggs and *T. latifolia* pollen to evaluate their nutritional value and suitability for rearing *T. montdorensis* under laboratory conditions. Overall, the application of frozen *C. lactis* eggs and *T. latifolia* pollen significantly reduced the immature development time and enhanced the reproductive capacity of *T. montdorensis*, compared to using living *C. lactis* as food for this predator. More importantly, this method can be applied to multiply several other Phytoseiidae species, thereby improving the effectiveness of biological control strategies across various crops.

## Materials and methods

### Stock colony of *T. montdorensis* and culture methods

The laboratory cultures of the predatory mites *T. montdorensis* were initially supplied by Bugs for Bugs (Queensland 4350, Australia) with food sources including mixed stages of living *C. lactis* and vermiculite as a shelter for the mites. Subsequently, this colony was maintained in a climatic set to 25 ± 1 °C, 70 ± 5 % RH, and a 16:8 h (L:D) photoperiod. The mites were maintained on white tiles (200 × 100 × 7 mm) (Johnson Tiles, Victoria 3153, Australia) that were placed on top of wet, thick foam pads (200 × 80 × 100 mm) (QEP company, Victoria 3175, Australia) inside a plastic tray (40 × 32 × 17 cm) filled with water. To prevent the mites from escaping and to ensure optimal humidity for predator development, the edges of the tiles were surrounded by tissue papers soaked in water. A curved piece of black filter paper (Westlab Pty Ltd, Victoria 3355, Australia) was placed in the centre of the tile to serve as shelter for the mites (Fig. 1a). Mixed stages of living *C. lactis* were added as food when needed.

**Fig. 1.**
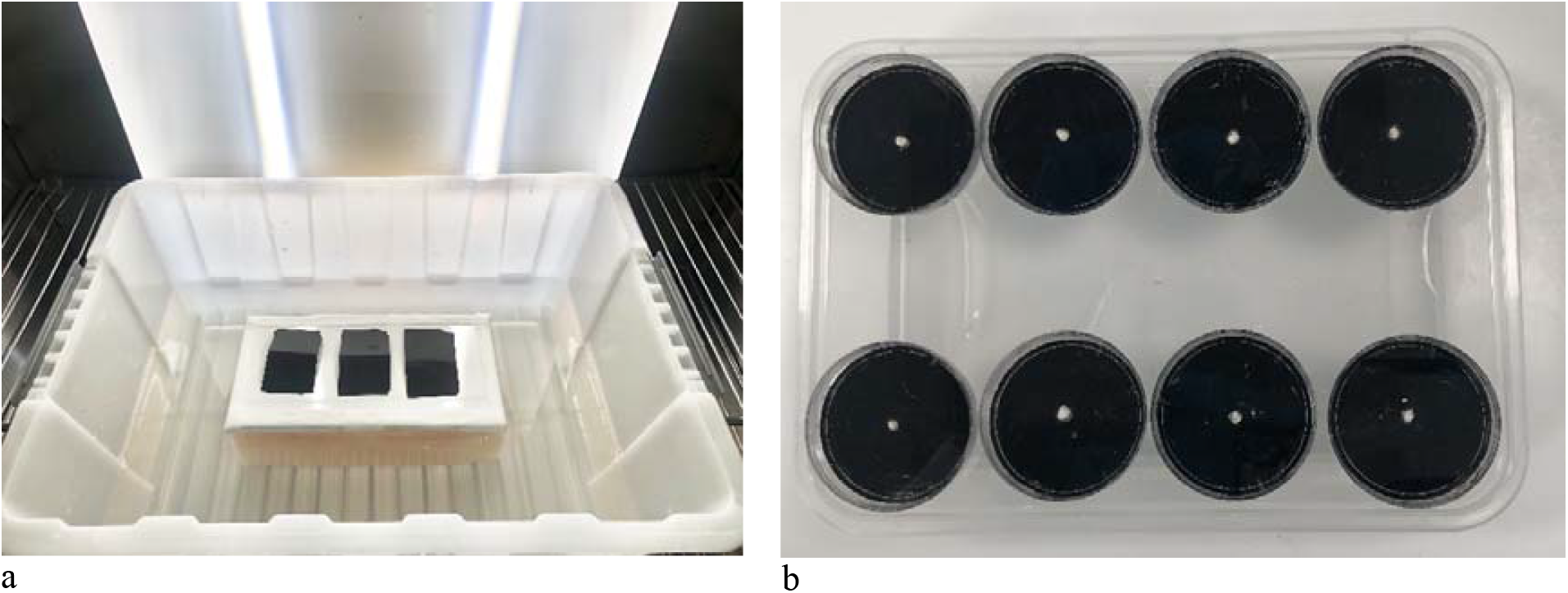
Rearing arenas of *Transeius montdorensis*. a. Stock colony, b. Rearing microcosms

### Diet preparation

#### Pollen

The fresh cattail pollen (*T. latifolia*) was collected in the cattail field in Menangle, New South Wales 2568, Australia, in December 2022 by tapping flowers on a tray and then transferred into a 50 ml tube (Thermo Fisher Scientific Australia Pty Ltd, Victoria 3179, Australia) before freezing at -20 °C. Prior to use in the experiments, the pollen was thawed and then stored at a temperature of 4 °C for up to one week.

#### Living C. lactis

A colony of *C. lactis* was also supplied by the Bugs for Bugs. The storage mites were reared in Petri dishes (90 × 15 mm). The prey mites were fed exclusively on instant dry yeast (Lowan Australia Ltd, NSW 2761, Australia) to collect frozen eggs of *C. lactis* by sieving through a fine net mesh. The Petri dishes, containing living *C. lactis* and its foods, were sealed with cling wrap (MPM Marketing Services, QLD 4008, Australia) and then placed on tiles (200 × 100 × 7 mm) (Johnson Tiles, Victoria 3153, Australia). These tiles were positioned on thick sponge pads inside a plastic box (40 × 32 × 17 cm) filled with water. A sharp needle was used to puncture tiny holes in the cling wrap to ensure ventilation within the Petri dishes while preventing prey escape. The stocks were kept at 25 ± 2 °C, 85 ± 5 % RH, and in complete darkness (Bakr et al., 2021) by covering the lids and all sides of the rearing containers with thick black plastic bags. The mixed life stages of *C. lactis* served as the food source for one colony of *T. montdorensis* and were used in experiments.

#### Frozen C. lactis eggs

To collect the frozen eggs of *C. lactis*, entire Petri dishes containing all life stages of the prey mites, as well as their food diets and feces, were frozen at -20 °C for at least 48 hours to ensure complete mortality. Subsequently, the contents were sieved through a fine mesh net (100µm) (Pathtech, Victoria 3072, Australia) to collect only *C. lactis* eggs. These eggs were then stored at - 20 °C for no longer than 4 weeks to ensure their quality (Pirayeshfar et al., 2020).

### Experimental design and statistical analyses

#### Stock colony of predatory mites T. montdorensis

There were four stocks of *T. montdorensis*, each fed on one of four different food diets: cattail pollen (*T. latifolia*), mixed stages of living *C. lactis*, frozen *C. lactis* eggs, and a combination of *T. latifolia* pollen and frozen *C. lactis* eggs in a 1:1 ratio (mixed diet). The rearing setup was similar to the main stock, except the white tiles were replaced by blue ones (97 × 97 mm). This alteration aimed to enable the easy detection and differentiation of male and female predators. Food was replenished every two days, with the removal of mouldy residues as necessary to prevent adverse effects on mite population development. Each population was reared for at least 2 generations before collecting fresh eggs for experimental setups.

#### Rearing microcosms

Small Petri dishes (35 × 12 mm) were used to examine the development and reproduction of individual *T. montdorensis*. To improve their development’s visibility, each Petri dish’s exterior was coated with black acrylic paint (Shamrock company, VIC 3130, Australia). Each dish featured a small hole at the bottom, into which a piece of cotton thread was inserted. A total of eight small Petri dishes were arranged within a single takeaway box equipped with a lid, which had eight corresponding holes, allowing the cotton threads to extend to the bottom of the box filled with tap water. This setup provided a water source to support the development of the predators. The Petri dishes were covered with cling wrap to prevent the mites from escaping while still allowing easy observation of their development and reproduction (Fig. 1b).

#### Experimental setup

To examine the development and reproduction of individual *T. montdorensis*, freshly laid eggs were collected within 8 hours and transferred each one into a separate small Petri dish for individual observation (n=36 for *T. latifolia* pollen and living *C. lactis* diets and n=32 for frozen *C. lactis* eggs and mixed diets). Upon egg hatching, the corresponding diets of *T. latifolia* pollen, living *C. lactis*, frozen *C. lactis* eggs, and mixed diet were introduced in the rearing microcosms. All food diets were replenished every two days except for the second diet, where living *C. lactis* in mixed stages were added as necessary. Observations were conducted every 12 hours until the mites reached the adult stage to collect data on the duration of each developmental stage, including egg and immature stages (the period after the egg hatched until the mite reached adulthood), as well as their survival rate. Developmental stages were determined based on the presence of cast skin in the rearing microcosms (Steiner et al., 2003). After emergence, each female was paired with a male fed on the same diet. Males that died during the experiment were replaced by other males that had been reared on the same diet. Subsequently, daily observations were made to assess oviposition periods, female longevity, and fecundity, while the preoviposition period was examined every 12 hours. To obtain the data on offspring’s sex ratio and survival proportion, eggs laid were collected daily, transferred to new rearing microcosms, and provided with the same diets as their parents until they reached adulthood. Mites that died as a result of non-natural causes, including accidental fatalities while attempting to recover the cling wrap on the rearing microcosm or escape from the rearing arena, were excluded from data analysis. The experiments were conducted in a growth chamber set to a constant environment of 25 ± 1 °C, 70 ± 5 % RH, and a 16:8 hours (L:D) photoperiod.

#### Statistical analysis

Statistical analysis was conducted using R software version 4.4.0 to evaluate the impact of diet on various aspects of the predatory mite’s biology, including the development time of egg and immature stages, preoviposition and oviposition duration, daily and total eggs laid, proportion of female offspring and female longevity. Initially, the Shapiro-Wilk test was performed to assess data normality. For non-normally distributed data, means were calculated using the “dplyr” package and compared using the “dunn.test” package. Dunn’s test with Bonferroni correction was used in conjunction with the Kruskal-Wallis test to calculate the chi-square (χ2), degrees of freedom (df), and P-value. To analyse the proportion of female offspring across four treatments and to compare their means, a Generalized Linear Model (GLM) was employed. This was followed by post-hoc analysis using Dunn’s test with Bonferroni correction to determine statistical significance and assign significance labels. Pairwise comparisons were made to identify differences between treatments, with a conventional alpha level of 0.05 used to assess significance. Therefore, P-values of 0.05 or lower were considered significant.

#### Life table parameters

The life table parameters, including the net productive rate (R_0_), generation time (Tc), and intrinsic rate of increase (r_m_), were calculated employing the Jackknife procedure as described by Maia et al. (2000).

## Results

### Developmental time

The survival proportion of individuals at the immature stages across all four treatments was 100%. For females, the total immature developmental time was shortest when fed on the mixed diet (5.30 ± 0.14 days) and longest for those reared on living *C. lactis* (6.05 ± 0.21 days). Notably, the developmental time of female deutonymphs fed on the mixed diet (1.15 ± 0.05 days) was significantly shorter compared to those reared on *T. latifolia* pollen (1.43 ± 0.08 days), living *C. lactis* (1.79 ± 0.15 days), and frozen *C. lactis* eggs (1.55 ± 0.08 days). There were no significant differences in the developmental time of egg, larval, and protonymph stages for females across the various diets. For males, the shortest total developmental time was observed with frozen *C. lactis* eggs (5.00 ± 0.17 days), which was significantly different from that recorded with cattail pollen (5.54 ± 0.14 days) and living *C. lactis* (5.97 ± 0.09 days), but not significantly different from the mixed diet (5.38 ± 0.20 days). Notably, the developmental time of the eggs that hatched into males, laid by females fed on frozen *C. lactis* eggs (1.65 ± 0.19 days) was significantly shorter than that of eggs from females reared on *T. latifolia* pollen (2.19 ± 0.17 days) and living *C. lactis* (2.18 ± 0.07 days). Regarding the larval stage, the developmental time of males fed on *T. latifolia* pollen (0.62 ± 0.06 days) was significantly shorter than those fed on the mixed diet (0.71 ±

0.07 days), living *C. lactis* (0.88 ± 0.05 days), and frozen *C. lactis* eggs (0.96 ± 0.09 days). However, the developmental time of the protonymph stage did not differ among the four diets. Meanwhile, male deutonymphs developed significantly faster when fed on frozen *C. lactis* eggs (1.15 ± 0.07 days) compared to those fed on *T. latifolia* pollen (1.38 ± 0.08), the mixture (1.38 ± 0.09 days), and living *C. lactis* (1.53 ± 0.08 days).

### Reproduction

Diet significantly influenced the reproductive parameters of the predatory mite *T. montdorensis*, including oviposition period, proportion of female offspring, daily oviposition rate, fecundity, and survival proportion of offspring (Table 2). Preoviposition was shortest in females fed on the mixed diet (2.08 ± 0.11 days), followed by frozen *C. lactis* eggs (2.28 ± 0.12 days), *T. latifolia* (2.61 ± 0.12 days), and living *C. lactis* (2.63 ± 0.17 days). In contrast, no significant differences were observed in the oviposition period and female longevity across the four diets, with duration ranging from 21.4 ± 1.12 days to 23.7 ± 1.62 days for the oviposition period and from 26.8 ± 1.22 days to 30.9 ± 1.83 days for female longevity. Fecundity was highest in females reared on the mixed diet (43.1 ± 2.07 eggs), followed by those on frozen *C. lactis* eggs (33.2 ± 2.10 eggs) and on *T. latifolia* pollen (29.6 ± 1.55 eggs) while the lowest fecundity was observed in females fed on living *C. lactis* (27.3 ± 1.38 eggs). The daily oviposition rate followed a similar trend, with the highest rate recorded in the mixed diet (1.59 ± 0.07 eggs/female/day) and the lowest one observed in living *C. lactis* (1.00 ± 0.06 eggs/female/day). Peak oviposition occurred on days 6, 8, 10, and 12 after emergence for diets of frozen *C. lactis* eggs, the mixed diet, *T. latifolia* pollen, and living *C. lactis*, respectively (Fig. 2). The proportion of female progeny was significantly higher in offspring reared on diets of *T. latifolia* pollen and the mixed diet (0.65 ± 0.03 %) compared to those on frozen *C. lactis* eggs (0.59 ± 0.04 %) and living *C. lactis* diets (0.46 ± 0.05 %).

**Table 1.**
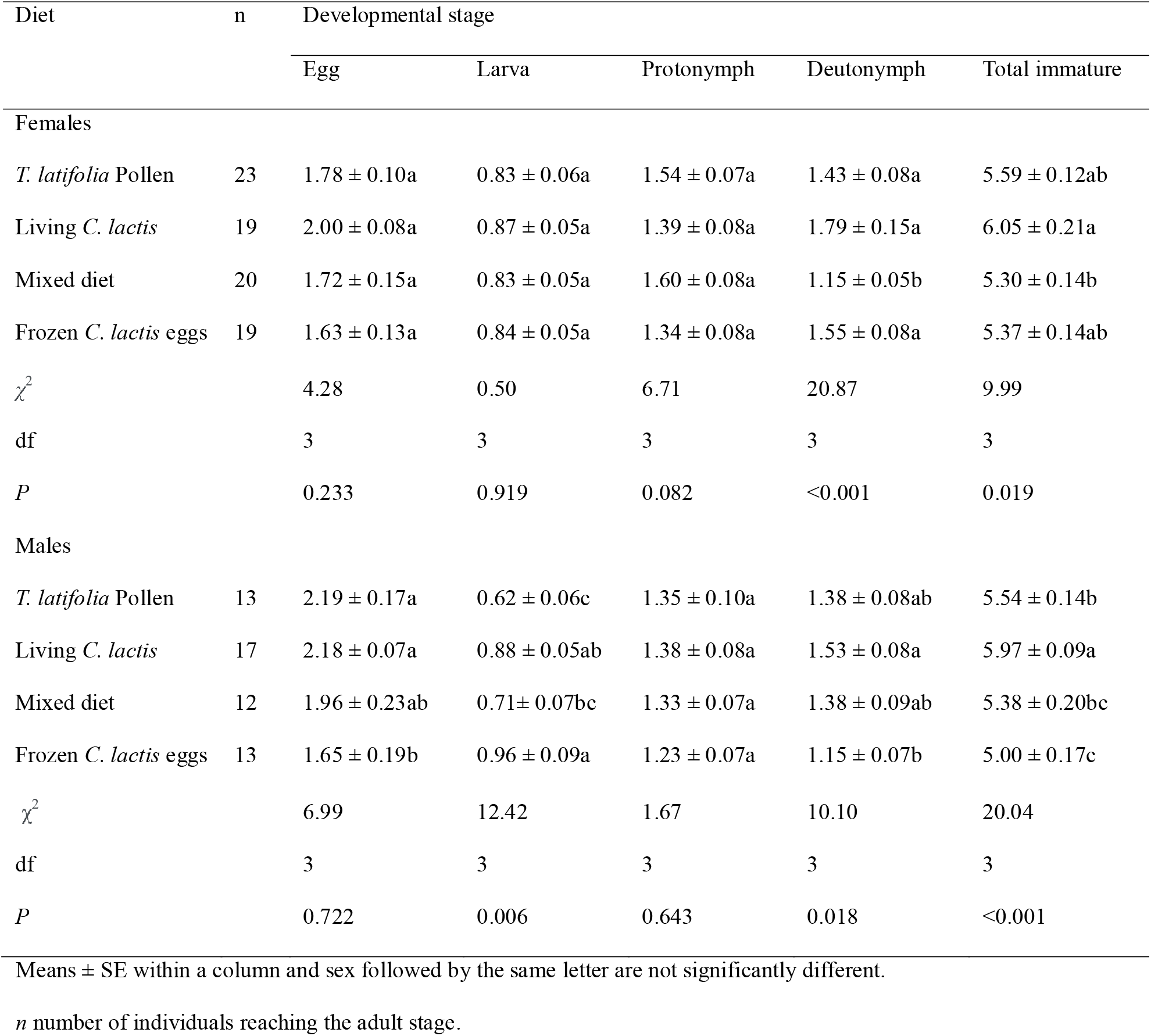
Developmental time (days) of the egg and immature stages of *Transeius montdorensis* fed on four diets at 25 °C.

**Table 2.**
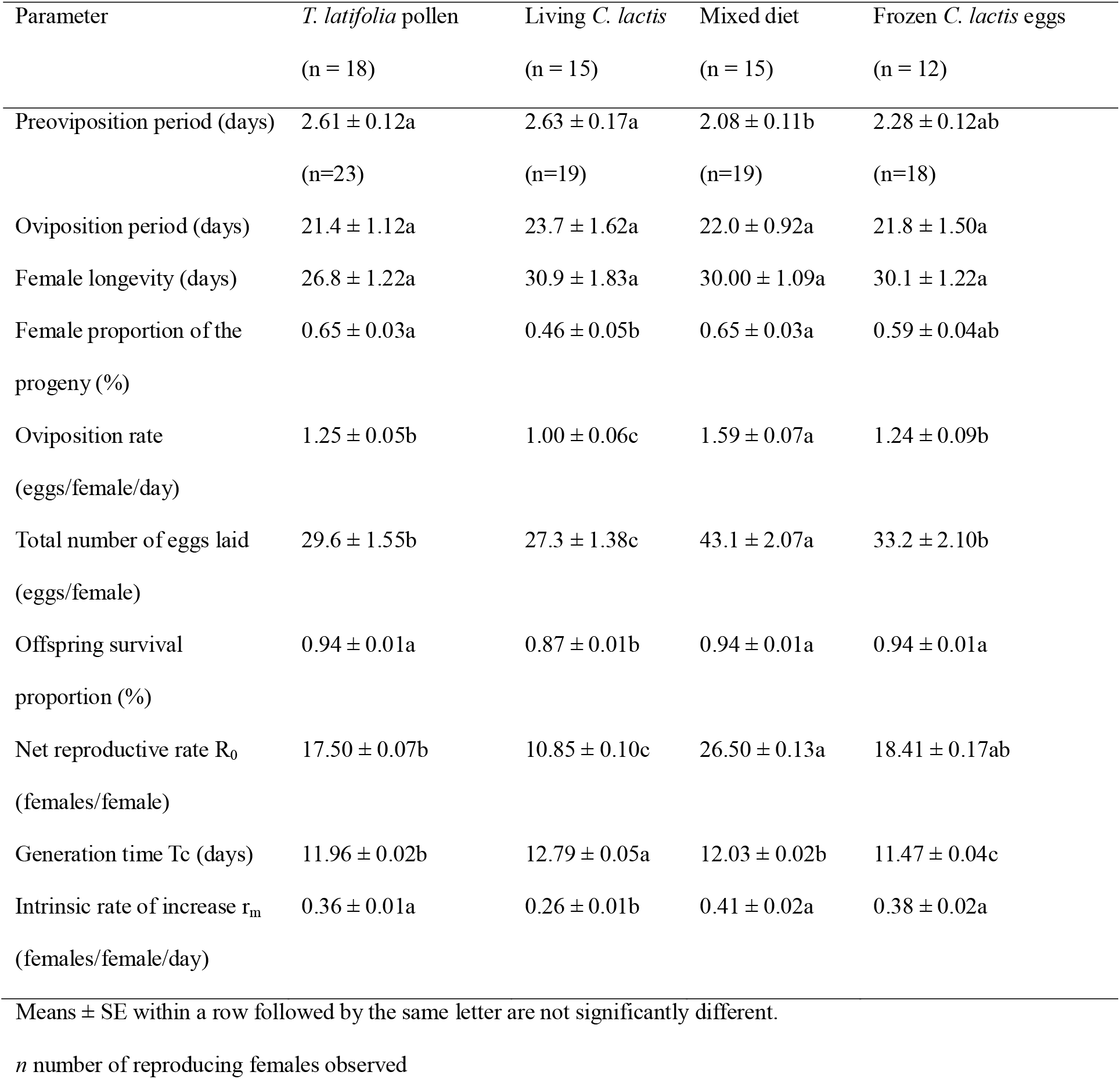
Reproduction and life table parameters of *Transeius montdorensis* fed on four diets at 25 °C.

**Fig. 2.**
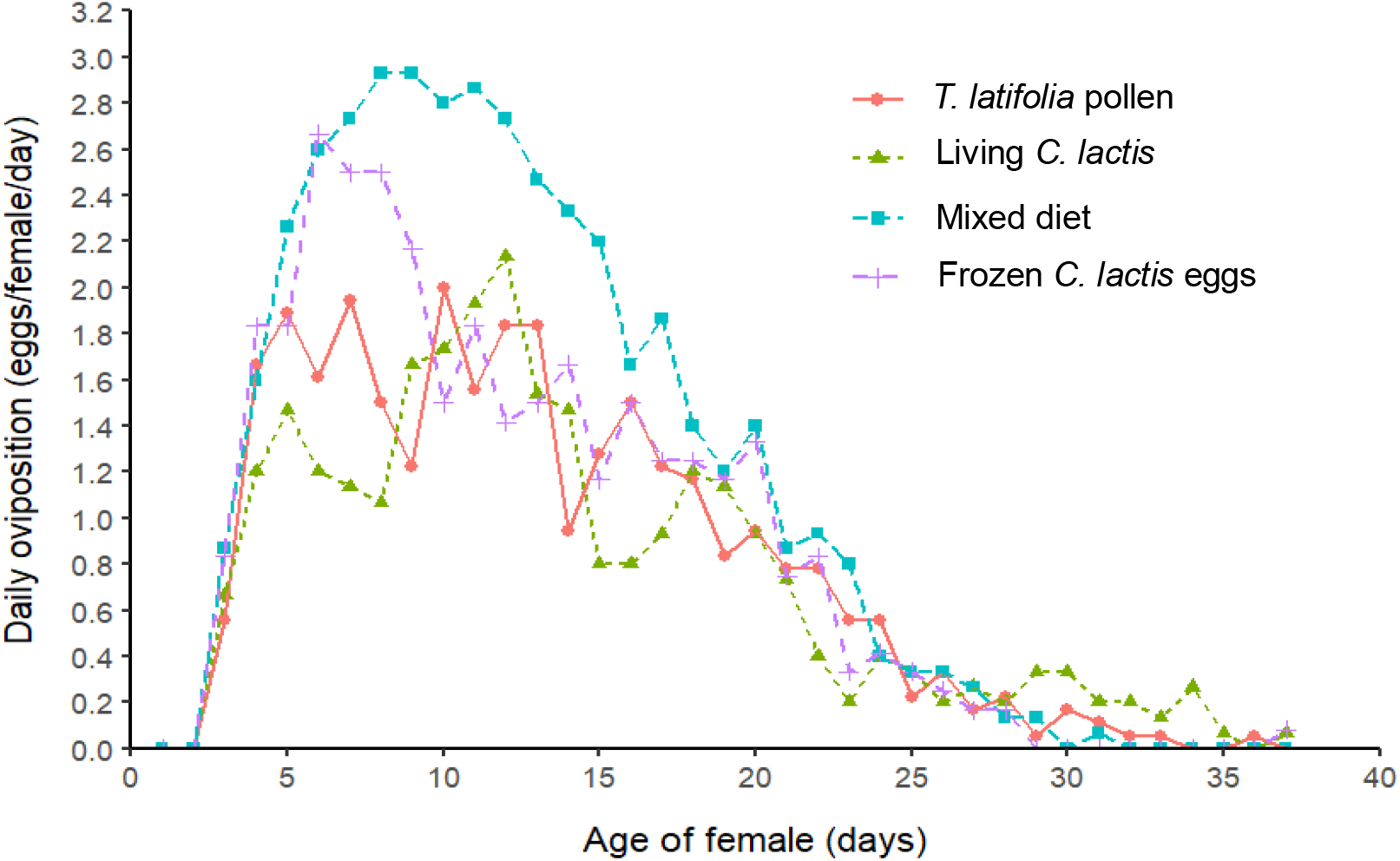
Daily oviposition rate of *Transeius montdorensis* fed on four diets at 25 °C

### Life table parameters

The net reproductive rate (R_0_) of *T. montdorensis* females that fed on the mixed diet (26.50 ± 0.13 females/female) was significantly higher than that of mites fed on *T. latifolia* pollen (17.50 ± 0.07 females/female) and living *C. lactis* (10.85 ± 0.10 females/female) (Table 2). The generation time of females fed on living *C. lactis* (12.79 ± 0.05 days) was significantly longer than those reared on the mixed diet (12.03 ± 0.02 days), *T. latifolia* pollen (11.96 ± 0.02 days). Meanwhile, females *T. montdorensis* that consumed frozen *C. lactis* eggs had the shortest generation period (11.47 ± 0.04 days). The intrinsic rate of increase (r_m_) for the females reared on the mixed diet (0.41 ± 0.02 female/female/day) was higher than those reared on frozen *C. lactis* eggs (0.38 ± 0.02 female/female/day), and those fed on *T. latifolia* pollen (0.36 ± 0.01 female/female/day), and significantly higher compared to those fed on living *C. lactis* (0.26 ± 0.01 female/female/day).

## Discussion

The primary objective of our research was to optimize the rearing method for the predatory mite *Transeius montdorensis* under controlled laboratory conditions. Our findings demonstrated a significant enhancement in development and reproduction for *T. montdorensis* females when fed on a combination of *T. latifolia* pollen and frozen *C. lactis* eggs. Particularly, the mixed diet reduced the total immature developmental time to 5.30 ± 0.14 days, notably shorter than that of females reared on other diets, especially living *C. lactis*, where the total development time of females was longest, at 6.05 ± 0.21 days. On the contrary, another study reported an immature development time of 7.5 ± 0.16 days for *T. montdorensis* fed on a combination of *Tetranychus urticae* and cattail pollen at 25 °C (Hatherly et al., 2004). Our study further highlights the value of the mixed diet by demonstrating superior life table parameters. Specifically, the highest value of both intrinsic rates of increase (r_m_) and net reproductive rate (R_0_) in *T. montdorensis* females reared on the combination of frozen *C. lactis* eggs and *T. latifolia* pollen. Females fed exclusively on cattail pollen exhibited a lower intrinsic rate of increase (r_m_ = 0.36) than those fed on the mixed diet (rm = 0.41). However, it was higher than reported by Steiner et al. (2003) under the same laboratory conditions (25 °C, 70 % RH, and a 16:8 h (L:D) photoperiod) (r_m_ = 0.32). This indicated that the mixed diet is preferable for *T. montdorensis*, as superior life table parameters result from dietary preference in arthropods, which subsequently leads to a high population build-up (Grenier & Clercq, 2003). Furthermore, the total number of eggs laid per female reared on the mixed diet was highest, averaging 43.1 ± 2.07 eggs per female, and almost double compared to that by females fed on living *C. lactis*, averaging 27.3 ± 1.38 eggs per female.

Pollen has been recognized as a promising alternative food source during periods of prey scarcity (He et al., 2022) and as a dietary supplement for large-scale rearing practices (Eini et al., 2022; I. Goleva & C. P. W. Zebitz, 2013; Yazdanpanah et al., 2021a). Despite recommendation by Hulshof and Vanninen (2002) to reduce pollen availability to manage *F. occidentalis*, subsequent research has shown that pollen positively affected the increase of predatory mite populations such as *A. limonicus* (Lee & Zhang, 2018; Samaras et al., 2019), *A. swirskii* (Barzka et al., 2023), and *N. cucumeris* (Han et al., 2024). Some species, for example, *Amblyseius cucumeris, Typhlodromus pyri*, and *Amblyseius andersoni* have been found to achieve comparable or even superior reproductive outcomes when fed on pollen rather than on their natural prey (Duso & Camporese, 1991; Marisa & Sauro, 1990). In a separate study, the supplementary of corn pollen to feed *Euseius scutalis* and *A. swirskii* resulted in a tenfold and twofold increase in their populations, respectively (Adar et al., 2014). Additionally, Gravandian et al. (2022) underscored the utility of *T. latifolia* pollen in sustaining immature development and fecundity of *N. cucumeris* females over 25 consecutive generations. However, pollen lacks β-carotene (Hatherly et al., 2004), which is rich in yeast, the primary food of *C. lactis*. Additional studies have also indicated the advantages of incorporating pollen into the diets of predatory mites under laboratory conditions (Eini et al., 2022; Irina Goleva & Claus P. Zebitz, 2013; Yazdanpanah et al., 2021b). Therefore, a combination of pollen and a protein-rich diet is necessary to improve the development and reproduction of predatory mites. This aligns with the results of this study, where life table parameter values, including R_0_, r_m_, and fecundity, for *T. montdorensis* females fed on the mixed diet were significantly higher than those fed on pollen alone. These results support the hypothesis that a mixed diet can enhance growth rates and reproductive outcomes, suggesting a synergistic effect from combining pollen with frozen prey, potentially due to improved nutritional balance.

The utilization of the storage mite *C. lactis* as an alternative diet offers several advantages over natural prey species. Studies have shown that *C. lactis* can be easily reared on simple media like yeast and sugar (Barbosa & de Moraes, 2015; San et al., 2020). Additionally, the high reproduction and rapid population growth of *C. lactis* ensures a consistent supply of food resources for the large-scale production of predatory mites. Consequently, *C. lactis* has been employed as a promising alternative food resource for rearing various predatory mites, including *T. montdorensis* and *A. swirskii* (Pirayeshfar et al., 2020; Zhang & Zhang, 2021). However, this study’s outcomes showed the lowest effectiveness of living *C. lactis* in both the development and reproduction of the predatory mites. *T. montdorensis* females fed on living *C. lactis* had the longest immature duration but the lowest net reproductive rate (R_0_), intrinsic rate of increase (r_m_), and total number of eggs laid. Moreover, using *C. lactis* as factitious prey is not without any drawbacks. Occasional observations during the experiments revealed that mature *C. lactis* occasionally preyed on the eggs and larvae of *T. montdorensis*, which likely contributed to the reduced fecundity observed in the predatory mites fed exclusively on this diet. Jan Hubert (2011) highlighted that the risk of *C. lactis* contamination has increased in the Czech Republic because *C. lactis* can migrate between packages in supermarkets. Additionally, contact with storage mites or even agricultural products that are infested with the mites can cause itching and redness or trigger storage-mite allergies in humans. The application of frozen *C. lactis* eggs can prevent the contamination risks associated with living *C. lactis*, thereby reducing the environmental impact when releasing *T. montdorensis* and frozen *C. lactis* eggs into greenhouse or field, rather than a combination involving living *C. lactis*.

To our knowledge, this study is the first to investigate the application of frozen *C. lactis* eggs in rearing the predatory mite *T. montdorensis*. Prior research on the efficacy of frozen eggs and the early stages of storage mites in supporting the development and fecundity of predators is limited. Contrary to our findings, such studies generally indicated that the performance of these alternative diets did not surpass those of living prey. Li and Zhang (2016) reported an extended juvenile development time for *N. cucumeris* when consuming frozen eggs of *T. urticae* as opposed to fresh eggs. In addition, research conducted by Xu et al. (2023) on the influence of frozen and fresh *T. urticae* eggs on the immature development, consumption rates, and oviposition rates of *Phytoseiulus persimilis* under laboratory conditions suggested that the nutritional quality of frozen eggs might be inadequate, leading to reduced oviposition rates and increased consumption rates when females are fed frozen prey mite eggs. Similarly, a study examining the impact of frozen eggs of storage mites *Tyrophgus putrescentiae* on the development and reproduction of *A. swirkii* illustrated that frozen *T. putrescentiae* eggs with cattail pollen did not significantly enhance the daily oviposition rate of this predator. The contrasting results regarding the effectiveness of frozen storage mite eggs for the development and reproduction of their predators between our study and others may be due to differences in prey and predator species, experimental setups, and methodology of each research (Pirayeshfar et al., 2020). Fatemeh Pirayeshfar et al. (2021) also reported that females of *B. mali* fed on one-day frozen eggs of storage mite *T. putrescentiae* exhibited higher life table parameters (r_m_, R_0,_ and λ) than those fed on eggs frozen for 90 days. It is, therefore, crucial to evaluate the developmental and reproductive performance of *T. montdorensis* females reared on frozen *C. lactis* eggs stored for different durations. Assessing the optimal ratio of frozen *C. lactis* eggs and *T. latifolia* pollen is also essential. Following these trials, further experiments should be scheduled to evaluate the feasibility of applying this mixed diet in mass-rearing.

Our results highlight the superiority of a combination of frozen *C. lactis* eggs and *T. latifolia* pollen in rearing *T. montdorensis* under laboratory conditions. This approach may not only support an effective mass-rearing program but also mitigate concerns associated with contamination and infestation by living *C. lactis*, as reported in previous studies (Çobanoğlu, 2009; Hubert et al., 2015; Jan Hubert, 2011). However, the duration for which prey mites are frozen could influence the reproduction outcomes of predatory mites. Moreover, extending this study to other predatory mites that consume *C. lactis*, such as *A. swirskii* (Asgari et al., 2020; Nguyen et al., 2013; San et al., 2020), *Phytoseiulus persimilis* (Tabic et al., 2022), *N. cucumeris* (Ji et al., 2015), and *Neoseiulus californicus* (Sweelam & Nasreldin, 2023; Wang et al., 2024) could provide broader implications for biological control strategies across different agricultural contexts.

## Acknowledgment

We are grateful to Dr Duong Nguyen of the Elizabeth Macarthur Agricultural Institute, NSW Department of Primary Industries, for generously providing *T. latifolia* pollen. We would also like to thank Dr Duc Tung Nguyen of the Entomology Department, Vietnam National University of Agriculture, for his guidance in calculating the life table parameters of *T. montdorensis*.

## Author contributions

Conceptualization: Hung Nguyen, Maciej Maselko, Bishwo Mainali, Binh Nguyen; Methodology: Hung Nguyen, Maciej Maselko, Bishwo Mainali, Binh Nguyen; Formal analysis and investigation: Hung Nguyen; Writing-original draft preparation: Hung Nguyen; Writing-review and editing: Maciej Maselko, Hung Nguyen, Binh Nguyen, Bishwo Mainali; Funding acquisition: Maciej Maselko; Resources: Department of Agriculture, Fisheries and Forestry; Supervision: Maciej Maselko

## Declarations

### Informed consent

The study does not contain any person’s data. Hence, informed consent is not applicable.

### Ethics

Biosafety approval was granted by the Macquarie University Institutional Biosafety Committee.

## References

Adar, E., Inbar, M., Gal, S., Gan-Mor, S., & Palevsky, E. (2014). Pollen on-twine for food provisioning and oviposition of predatory mites in protected crops. BioControl, 59, 307–317.

Asgari, F., Moayeri, H. R. S., Kavousi, A., Enkegaard, A., & Chi, H. (2020). Demography and Mass Rearing of Amblyseius swirskii (Acari: Phytoseiidae) Fed on Two Species of Stored-Product Mites and Their Mixture. Journal of Economic Entomology, 113(6), 2604–2612.

Bakr, A., Rezk, H. A., El-Hamid, A., Madiha, M., & Osman, S. I. (2021). Studying the Biology of Carpoglyphus lactis (L.) Reared on Dried Apricots and Its Control Using Plant Oil Extracts. Alexandria Science Exchange Journal, 42(2), 407–411.

Barbosa, M. F., & de Moraes, G. J. (2015). Evaluation of astigmatid mites as factitious food for rearing four predaceous phytoseiid mites (Acari: Astigmatina; Phytoseiidae). Biological Control, 91, 22–26.

Barzka, M., Shishehbo, P., Habibpou, B., Hemmat, A., & Riah, E. (2023). Development, survival, and reproduction of Amblyseius swirskii (Athias-Henriot)(Acari: Phytoseiidae) feeding on different pollen grains. Acarologia, 63(4), 1062–1071.

Bazgir, F., Shakarami, J., & Jafari, S. (2019). Life table and predation rate of Typhlodromus bagdasarjani (Acari: Phytoseiidae) fed on Eotetranychus frosti (Tetranychidae) on apple leaves. INTERNATIONAL JOURNAL OF ACAROLOGY, 45(4), 202–208.

Buitenhuis, R., Shipp, L., & Scott-Dupree, C. (2010). Intra-guild vs extra-guild prey: effect on predator fitness and preference of Amblyseius swirskii (Athias-Henriot) and Neoseiulus cucumeris (Oudemans)(Acari: Phytoseiidae). Bulletin of entomological research, 100(2), 167–173.

Chant, D. (1985). Systematics and morphology. Spider mites: their biology, natural enemies and control, 1, 3–10.

Çobanoğlu, S. (2009). Mite population density analysis of stored dried apricots in Turkey. INTERNATIONAL JOURNAL OF ACAROLOGY, 35(1), 67–75.

Cox, P., Matthews, L., Jacobson, R., Cannon, R., MacLeod, A., & Walters, K. (2006). Potential for the use of biological agents for the control of Thrips palmi (Thysanoptera: Thripidae) outbreaks. Biocontrol Science and Technology, 16(9), 871–891.

Cuthbertson, A. G. (2014). The feeding rate of predatory mites on life stages of Bemisia tabaci Mediterranean species. Insects, 5(3), 609–614.

Demite, P. R., McMurtry, J. A., & De Moraes, G. J. (2014). Phytoseiidae Database: a website for taxonomic and distributional information on phytoseiid mites (Acari). Zootaxa, 3795(5), 571–577-571–577.

Duso, C., & Camporese, P. (1991). Developmental times and oviposition rates of predatory mites Typhlodromus pyri and Amblyseius andersoni (Acari: Phytoseiidae) reared on different foods. Experimental & Applied Acarology, 13(2), 117–128.

Eini, N., Jafari, S., Fathipour, Y., & Zalucki, M. P. (2022). How pollen grains of 23 plant species affect performance of the predatory mite Neoseiulus californicus. BioControl, 67(2), 173–187.

Etienne, L., Bresch, C., van Oudenhove, L., & Mailleret, L. (2021). Food and habitat supplementation promotes predatory mites and enhances pest control. Biological Control, 159, 104604.

Goleva, I., & Zebitz, C. P. (2013). Suitability of different pollen as alternative food for the predatory mite Amblyseius swirskii (Acari, Phytoseiidae). Experimental & Applied Acarology, 61(3), 259–283.

Goleva, I., & Zebitz, C. P. (2013). Suitability of different pollen as alternative food for the predatory mite Amblyseius swirskii (Acari, Phytoseiidae). Experimental and Applied Acarology, 61, 259–283.

Goleva, I., & Zebitz, C. P. W. (2013). Suitability of different pollen as alternative food for the predatory mite Amblyseius swirskii (Acari, Phytoseiidae). Experimental and Applied Acarology, 61(3), 259–283.

Gravandian, M., Fathipour, Y., Hajiqanbar, H., Riahi, E., & Riddick, E. W. (2022). Long-term effects of cattail Typha latifolia pollen on development, reproduction, and predation capacity of Neoseiulus cucumeris, a predator of Tetranychus urticae. BioControl, 1–12.

Grenier, S., & Clercq, P. d. (2003). Comparison of artificially vs. Naturally reared natural enemies and their potential for use in biological control. CABI, 115–131.

Han, Y., Lipeizhong, W., Liang, X., Cai, Z., Liu, W., Dou, J., Lu, Y., Zhang, J., Wang, S., & Su, J. (2024). Effects of Various Nectar and Pollen Plants on the Survival, Reproduction, and Predation of Neoseiulus bicaudus. Insects, 15(3), 190.

Hatherly, I., Bale, J., Walters, K., & Worland, M. (2004). Thermal biology of Typhlodromips montdorensis: implications for its introduction as a glasshouse biological control agent in the UK. Entomologia experimentalis et applicata, 111(2), 97–109.

He, Y. M., Li, G. Y., Liu, M. X., Liu, H., & Wang, Z. Y. (2022). Effects of supplementary pollen on the life history traits of predatory mite Euseius nicholsi across generations. Journal of Applied Entomology, 146(10), 1293–1301.

Hubert, J., Nesvorna, M., & Volek, V. (2015). Stored product mites (Acari: Astigmata) infesting food in various types of packaging. Experimental and Applied Acarology, 65, 237–242.

Hubert, J., Stejskal, V., Athanassiou, C. G., & Throne, J. E. (2018). Health hazards associated with arthropod infestation of stored products. Annual review of entomology, 63(1), 553–573.

Hulshof, J., & Vanninen, I. (2002). Western flower thrips feeding on pollen, and its implications for control. sThrips and tospoviruses: proceedings of the 7th international symposium on Thysanoptera,

JA McMurtry, & Croft, B. (1997). Life-styles of Phytodeiid mites and their roles in biological control. In Annu. Rev. Entomol (pp. 291–321).

Jan Hubert, T. E., Marta Nesvorna & Vaclav Stejskal. (2011). Emerging risk of infestation and contamination of dried fruits by mites in the Czech Republic. Food Additives and Contaminants, 28(9), 7.

Jaques, J. A., Aguilar-Fenollosa, E., Hurtado-Ruiz, M. A., & Pina, T. (2015). Food web engineering to enhance biological control of Tetranychus urticae by phytoseiid mites (Tetranychidae: Phytoseiidae) in Citrus. Prospects for Biological control of plant feeding mites and other harmful organisms, 251–269.

Javier, F., Saenz-de-Cabezon, I., & Francisco, L.-O. J. (2010). A review of recent patents on macroorganisms as biological control agents. Recent Patents on Biotechnology, 4(1), 48–64.

Ji, J., Zhang, Y.-X., Lin, J.-Z., Chen, X., Sun, L., & Saito, Y. (2015). Life histories of three predatory mites feeding upon Carpoglyphus lactis (Acari, Phytoseiidae; Carpoglyphidae). Systematic and Applied Acarology, 20(5), 491–496.

Labbé, R. M., Gagnier, D., & Shipp, L. (2019). Comparison of Transeius montdorensis (Acari: Phytoseiidae) to Other Phytoseiid Mites for the Short-Season Suppression of Western Flower Thrips, Frankliniella occidentalis (Thysanoptera: Thripidae). Environmental Entomology, 48(2), 335–342.

Lee, M. H., & Zhang, Z.-Q. (2018). Assessing the augmentation of Amblydromalus limonicus with the supplementation of pollen, thread, and substrates to combat greenhouse whitefly populations. Scientific Reports, 8(1), 12189.

Li, G.-Y., & Zhang, Z.-Q. (2016). Some factors affecting the development, survival and prey consumption of Neoseiulus cucumeris (Acari: Phytoseiidae) feeding on Tetranychus urticae eggs (Acari: Tetranychidae). Systematic and Applied Acarology, 21(5), 555–566.

Lundgren, J. G. (2009). Relationships of natural enemies and non-prey foods (Vol. 7). Springer Science & Business Media.

Maia, A. d. H., Luiz, A. J., & Campanhola, C. (2000). Statistical inference on associated fertility life table parameters using jackknife technique: computational aspects. Journal of Economic Entomology, 93(2), 511–518.

Manners, A. G., Dembowski, B. R., & Healey, M. A. (2013). Biological control of western flower thrips, F rankliniella occidentalis (P ergande)(T hysanoptera: T hripidae), in gerberas, chrysanthemums and roses. Australian Journal of Entomology, 52(3), 246–258.

Marisa, C., & Sauro, S. (1990). Biological observations and life table parameters of Amblyseius cucumeris (Oud.)(Acarina: Phytoseiidae) reared on different diets.

Massaro, M., Montrazi, M., Melo, J. W. S., & de Moraes, G. J. (2021). Small-scale production of Amblyseius tamatavensis with Thyreophagus cracentiseta (Acari: Phytoseiidae, Acaridae). Insects, 12(10), 848.

McMurtry, J. (2010). Concepts of classification of the Phytoseiidae: Relevance to biological control of mites. Trends in acarology: Proceedings of the 12th International Congress,

McMurtry, J. A., De Moraes, G. J., & Sourassou, N. F. (2013). Revision of the lifestyles of phytoseiid mites (Acari: Phytoseiidae) and implications for biological control strategies. Systematic and Applied Acarology, 18(4), 297–320.

Mouratidis, A., Marrero-Díaz, E., Sánchez-Álvarez, B., Hernández-Suárez, E., & Messelink, G. J. (2023). Preventive releases of phytoseiid and anthocorid predators provided with supplemental food successfully control Scirtothrips in strawberry. BioControl, 68(6), 603–615.

Nguyen, D. T. (2015). Artificial and factitious foods for the production and population enhancement of phytoseiid predatory mites. Ghent: Ghent University. Faculty of Bioscience Engineering.

Nguyen, D. T., Vangansbeke, D., Lü, X., & De Clercq, P. (2013). Development and reproduction of the predatory mite Amblyseius swirskii on artificial diets. BioControl, 58, 369–377.

Pirayeshfar, F., Moayeri, H. R. S., Da Silva, G. L., & Ueckermann, E. A. (2022). Comparison of biological characteristics of the predatory mite Blattisocius mali (Acari: Blattisocidae) reared on frozen eggs of Tyrophagus putrescentiae (Acari: Acaridae) alone and in combination with cattail and olive pollens. Systematic and Applied Acarology, 27(3), 399–409.

Pirayeshfar, F., Safavi, S. A., Sarraf-Moayeri, H. R., & Messelink, G. J. (2021a). Active and frozen host mite Tyrophagus putrescentiae (Acari: Acaridae) influence the mass production of the predatory mite Blattisocius mali (Acari: Blattisociidae): life table analysis. Systematic and Applied Acarology, 26(11), 2096-2108, 2013.

Pirayeshfar, F., Safavi, S. A., Sarraf-Moayeri, H. R., & Messelink, G. J. (2021b). Provision of astigmatid mites as supplementary food increases the density of the predatory mite Amblyseius swirskii in greenhouse crops, but does not support the omnivorous pest, western flower thrips. BioControl, 66(4), 511–522.

Pirayeshfar, F., Safavi, S. A., Sarraf Moayeri, H. R., & Messelink, G. J. (2020). The potential of highly nutritious frozen stages of Tyrophagus putrescentiae as a supplemental food source for the predatory mite Amblyseius swirskii. Biocontrol Science and Technology, 30(5), 403–417.

Rahman, T., Spafford, H., & Broughton, S. (2011). Compatibility of spinosad with predaceous mites (Acari) used to control Frankliniella occidentalis (Pergande)(Thysanoptera: Thripidae). Pest management science, 67(8), 993–1003.

Rahman, T., Spafford, H., & Broughton, S. (2012). Use of spinosad and predatory mites for the management of Frankliniella occidentalis in low tunne□grown strawberry. Entomologia experimentalis et applicata, 142(3), 258–270.

Ranabhat, N. B., Goleva, I., & Zebitz, C. P. (2014). Life tables of Neoseiulus cucumeris exclusively fed with seven different pollens. BioControl, 59, 195–203.

Richter, E. (2016). Efficacy of two predatory mite species to control whiteflies infesting poinsettia plants compared to the standard parasitoid Encarsia formosa. III International Symposium on Organic Greenhouse Horticulture 1164,

Samaras, K., Pappas, M. L., Fytas, E., & Broufas, G. D. (2015). Pollen suitability for the development and reproduction of Amblydromalus limonicus (Acari: Phytoseiidae). BioControl, 60, 773–782.

Samaras, K., Pappas, M. L., Fytas, E., & Broufas, G. D. (2019). Pollen provisioning enhances the performance of Amblydromalus limonicus on an unsuitable prey. Frontiers in Ecology and Evolution, 7, 122.

San, P. P., Tuda, M., Nakahira, K., & Takagi, M. (2020). Optimal rearing medium for the population growth of the predatory mite, Amblyseius swirskii (Athias-Henriot)(Acari: Phytoseiidae). Egyptian Journal of Biological Pest Control, 30, 1–5.

Schicha, E. (1979). Three new species of “Amblyseius” Berlese from New Caledonia and Australia (Acari: Phytoseiidae) [Journal Article]. Australian Entomologist, 6(3), [41]-48.

Schreiber, I. (2018). The role of pollen as alternative food for predatory mites (Acari: Phytoseiidae).

Steiner, M., & Goodwin, S. (2002). Development of a new thrips predator, Typhlodromips montdorensis (Schicha)(Acari: Phytoseiidae) indigenous to Australia.

Steiner, M. Y., Goodwin, S., Wellham, T. M., Barchia, I. M., & Spohr, L. J. (2003). Biological studies of the Australian predatory mite Typhlodromips montdorensis (Schicha)(Acari: Phytoseiidae), a potential biocontrol agent for western flower thrips, Frankliniella occidentalis (Pergande)(Thysanoptera: Thripidae). Australian Journal of Entomology, 42(2), 124–130.

Sun, L., Liao, Z.-X., Zheng, Y.-Q., Chen, D.-S., Gao, G.-G., & Chen, X. (2022). Effects of temperature on immature development of Transeius montdorensis (Schicha)(Acari: Phytoseiidae) fed on Bemisia tabaci Gennadius (Hemiptera, Aleyrodidae) biotype Q. Systematic and Applied Acarology, 27(10), 2004–2011.

Sweelam, M. E., & Nasreldin, M. (2023). Biological aspects and life-tables of the predatory mites, Amblyseius swirskii Athias-Henriot and Neoseiulus californicus (McGregor), reared on four types of food. Acarines: Journal of the Egyptian Society of Acarology, 17(1), 45–55.

Tabic, A., Katz, T., & Steinberg, S. (2022). A newly characterized Phytoseiulus persimilis is a component of a novel mass rearing method and a revolutionary slow-release product. Zoosymposia, 22, 259–259.

Van Lenteren, J. C. (2012). The state of commercial augmentative biological control: plenty of natural enemies, but a frustrating lack of uptake. BioControl, 57(1), 1–20.

Vangansbeke, D., Nguyen, D. T., Audenaert, J., Verhoeven, R., Gobin, B., Tirry, L., & De Clercq, P. (2014). Performance of the predatory mite Amblydromalus limonicus on factitious foods. BioControl, 59(1), 67–77.

Wang, J., Zhang, K., Li, L., & Zhang, Z.-Q. (2024). Development and reproduction of four predatory mites (Parasitiformes: Phytoseiidae) feeding on the spider mites Tetranychus evansi and T. urticae (Trombidiformes: Tetranychidae) and the dried fruit mite Carpoglyphus lactis (Sarcoptiformes: Carpoglyphidae). Systematic and Applied Acarology, 29(2), 269–284.

Xu, Y., Zhang, K., & Zhang, Z.-Q. (2023). Development, survival, and reproduction of Phytoseiulus persimilis Athias-Henriot (Acari: Phytoseiidae) feeding on fresh versus frozen eggs of Tetranychus urticae Koch (Acari: Tetranychidae). Acarologia, 63(1), 24–30.

Yazdanpanah, S., Fathipour, Y., & Riahi, E. (2021a). Pollen grains are suitable alternative food for rearing the commercially used predatory mite Neoseiulus cucumeris (Acari: Phytoseiidae). Systematic and Applied Acarology, 26(5), 1009-1020, 1012.

Yazdanpanah, S., Fathipour, Y., & Riahi, E. (2021b). Pollen grains are suitable alternative food for rearing the commercially used predatory mite Neoseiulus cucumeris (Acari: Phytoseiidae). Systematic and Applied Acarology, 26(5), 1009–1020.

Zhang, K., & Zhang, Z.-Q. (2021). The dried fruit mite Carpoglyphus lactis (Acari: Carpoglyphidae) is a suitable alternative prey for Amblyseius herbicolus (Acari: Phytoseiidae). Systematic and Applied Acarology, 26(11), 2167-2176, 2110.

